# Message in a Bottle – Metabarcoding Enables Biodiversity Comparisons Across Ecoregions

**DOI:** 10.1101/2021.07.05.451165

**Authors:** D Steinke, SL deWaard, JE Sones, NV Ivanova, SWJ Prosser, K Perez, TWA Braukmann, M Milton, EV Zakharov, JR deWaard, S Ratnasingham, PDN Hebert

## Abstract

**Background:** Traditional biomonitoring approaches have delivered a basic understanding of biodiversity, but they cannot support the large-scale assessments required to manage and protect entire ecosystems. This study employed DNA metabarcoding to assess spatial and temporal variation in species richness and diversity in arthropod communities from 52 protected areas spanning three Canadian ecoregions.

**Results:** This study revealed the presence of 26,263 arthropod species in the three ecoregions and indicated that at least another 3,000–5,000 await detection. Results further demonstrate that communities are more similar within than between ecoregions, even after controlling for geographical distance. Overall α-diversity declined from east to west, reflecting a gradient in habitat disturbance. Shifts in species composition were high at every site with turnover greater than nestedness, suggesting the presence of many transient species.

**Conclusions:** Differences in species composition among their arthropod communities confirm that ecoregions are a useful synoptic for biogeographic patterns and for structuring conservation efforts. The present results also demonstrate that metabarcoding enables large-scale monitoring of shifts in species composition, making it possible to move beyond the biomass measurements that have been the key metric employed in prior efforts to track change in arthropod communities.

## Background

Terrestrial organisms are exposed to diverse anthropogenic stressors, including climate change, resource extraction, and agriculture. Habitat degradation, pesticide usage, invasive species, and associated shifts in food webs have provoked major reductions in the abundance of terrestrial arthropods [1-4]. These declines have led to calls for more comprehensive biosurveillance to inform environmental management and conservation. Long-term monitoring of species composition is essential to quantify biological change, but efforts employing morphological diagnostics have targeted a small set of indicator species because of the need for taxonomic experts for each group. As a consequence, they cannot support the broad assessments needed to manage and protect ecosystems, let alone forecast human impacts on them by integrating statistical modelling. The latter methods demand comprehensive data on species distributions and abundance [5], information that is currently unavailable because of the prior focus on selected biotic compartments at limited geographic scale.

Two methodological advances promise to meet the need for comprehensive biodiversity data. Firstly, identification systems based on the analysis of sequence variation in short, standardized gene regions (i.e., DNA barcodes) enable species discrimination [6]. Secondly, high-throughput sequencers (HTS) permit the inexpensive acquisition of millions of DNA barcode records [7]. These advances now enable biodiversity surveys at speeds and scales that were previously inconceivable. In particular, the coupling of HTS with DNA barcoding, known as metabarcoding [8], has a compelling advantage over traditional approaches for tracking shifts in species presence. It can generate georeferenced occurrence data from bulk samples at low cost, and a single instrument can process hundreds of bulk samples each week. Because the sequencing output of HTS is doubling every nine months [9,10], analytical costs are certain to sharply decline, allowing production to soar. This augmented capacity for data generation has already enabled large-scale biotic surveys of aquatic and terrestrial arthropods [11-14], vertebrates [15], pollen [16], diatoms [17], and fungi [18-20].

Access to large collections of specimens is essential to capitalize on the analytical capacity provided by DNA metabarcoding. Among the many approaches used to sample terrestrial arthropods, Malaise traps [21] have gained wide adoption because they collect large, diverse samples with little effort [22]. Although most-effective for sampling flying insects, they also collect ground-active arthropods. By coupling DNA barcoding with Malaise trapping [23,24], high-resolution monitoring networks for arthropods are within reach, but there are challenges. Data interpretation requires a well-parameterized DNA barcode reference library for the region under investigation, creating the need for a system to aid site selection. Ecoregions represent an obvious candidate [25-28] although their boundaries are rarely sharply defined, and they are based on distributional data for a narrow range of taxa. Despite these limitations, ecoregions have been widely used to guide management decisions and to explore species and community diversity patterns [29,30]. As a result, they are a good candidate to serve as the backbone for a large-scale monitoring network. The most widely adopted schema partitions the world’s 14 terrestrial biomes into 846 ecoregions [30].

This study demonstrates the feasibility of employing metabarcoding for large-scale bio-surveillance by comparing the temporal and spatial patterning of arthropod communities in three of Canada’s 47 terrestrial ecoregions: the Eastern Canadian Forest – Boreal Transition (ECF – 75,000 km^2^), the Eastern Great Lakes Lowland Forests (EGL – 63,000 km^2^), and the Southern Great Lakes Forests (SGL – 22,000 km^2^) (Figure 1). Forest cover declines from 77.7% in the ECF to 30.1% in the EGL and just 12.1% in the SGL while cropland/pastures cover 78% of the SGL, 57% of the EGL, and 3% of the ECF [31]. The EGL and SGL are the most populated ecoregions in Ontario with developed land (e.g., urban, road networks) encompassing more than 7% of the SGL [31]. As such, these ecoregions provide a good basis for assessing the impacts of varied disturbance regimes on biodiversity.

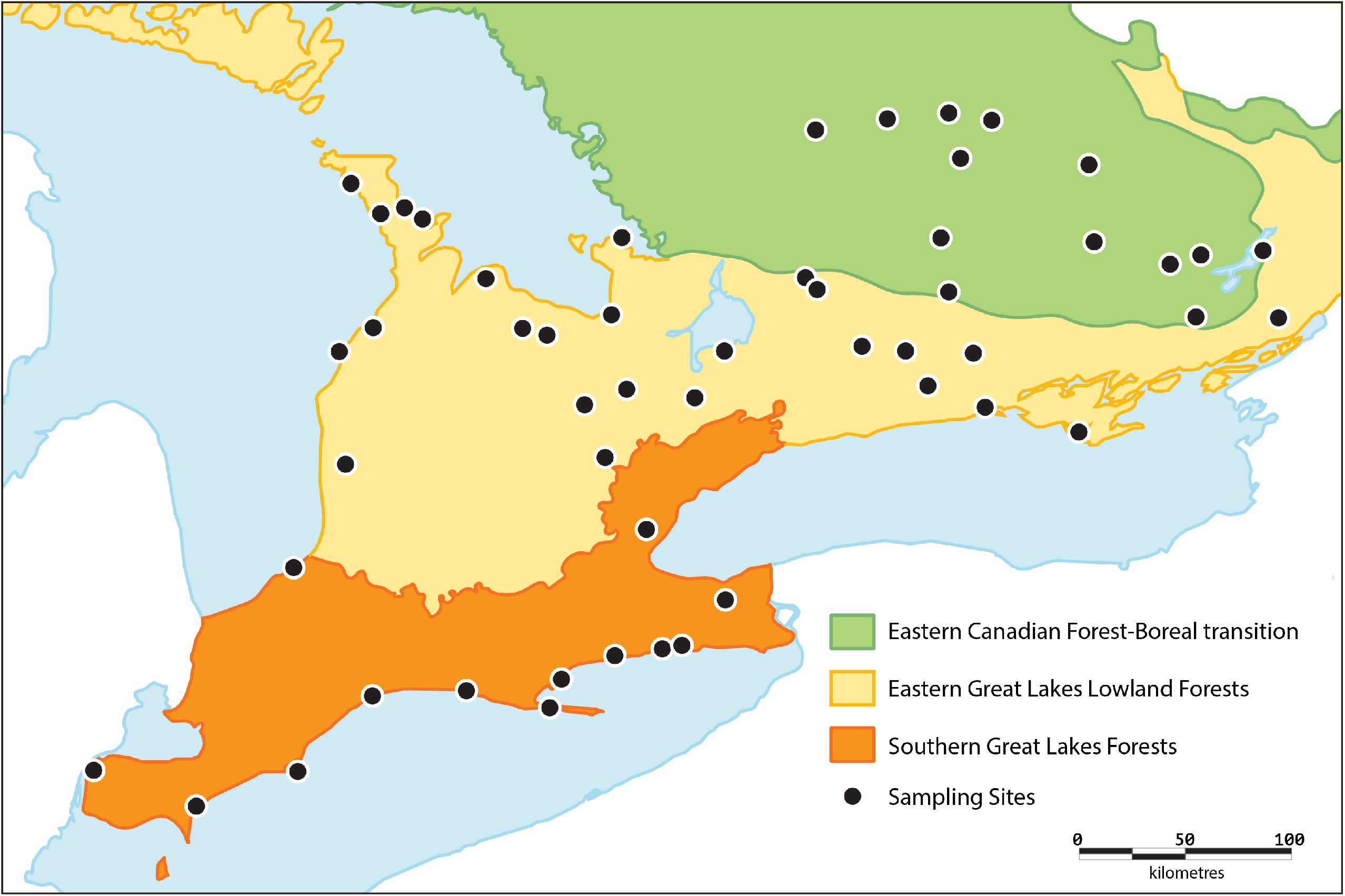

## Data Description

Collections were made by deploying a Malaise trap at 52 sites in these three ecoregions and samples were metabarcoded to examine variation in their species richness, community composition, phylogenetic diversity, as well as alpha (α) and beta (β)-diversity. Malaise traps were deployed for 20 weeks at 15 sites in the ECF, 24 sites in the EGL, and 13 sites in the SGL. Catches were harvested at two-week intervals and 410 of the resultant 520 samples were designated for metabarcoding (the others were reserved for single specimen barcoding). Analysis began with non-destructive lysis of the specimens in each bi-weekly sample, followed by DNA extraction using a membrane-based protocol [32]. A 463 bp amplicon of cytochrome *c* oxidase I (COI) was then PCR amplified and the amplicon pools from each set of 10 samples were sequenced on an Ion Torrent S5 using a 530 chip. The sequences were subsequently analyzed using the Multiplex Barcode Research And Visualization Environment (mBRAVE – mbrave.net). All raw HTS datasets were deposited in the Sequence Read Archive (SRA – www.ncbi.nlm.nih.gov/sra/) under the BioProject accession number PRJNA629553.

## Analyses

Sequence analysis of the 410 samples produced 367,823,207 reads across 41 S5 runs (mean reads per run = 8.97 million, see **Table S1**). Two thirds were filtered, leaving 126,253,260 reads that could be assigned to a BIN (Barcode Index Number; [33]) on BOLD [34] (**Figure S1**). Nearly all reads (99.3 %) found a BIN match on BOLD, but those that failed were *de novo* clustered using mBRAVE with a 99% similarity threshold. The latter analysis recognized an average of 28 additional OTUs per sample, but >96% of them reflected sequencing/PCR errors (e.g., chimeras, sequences with multiple indels) or NUMTs so they were excluded from further analysis. Consideration of the assigned reads revealed 26,263 BINs among the 52 sites with more than a third (9,301) found at only one site (**Figure 2b**).

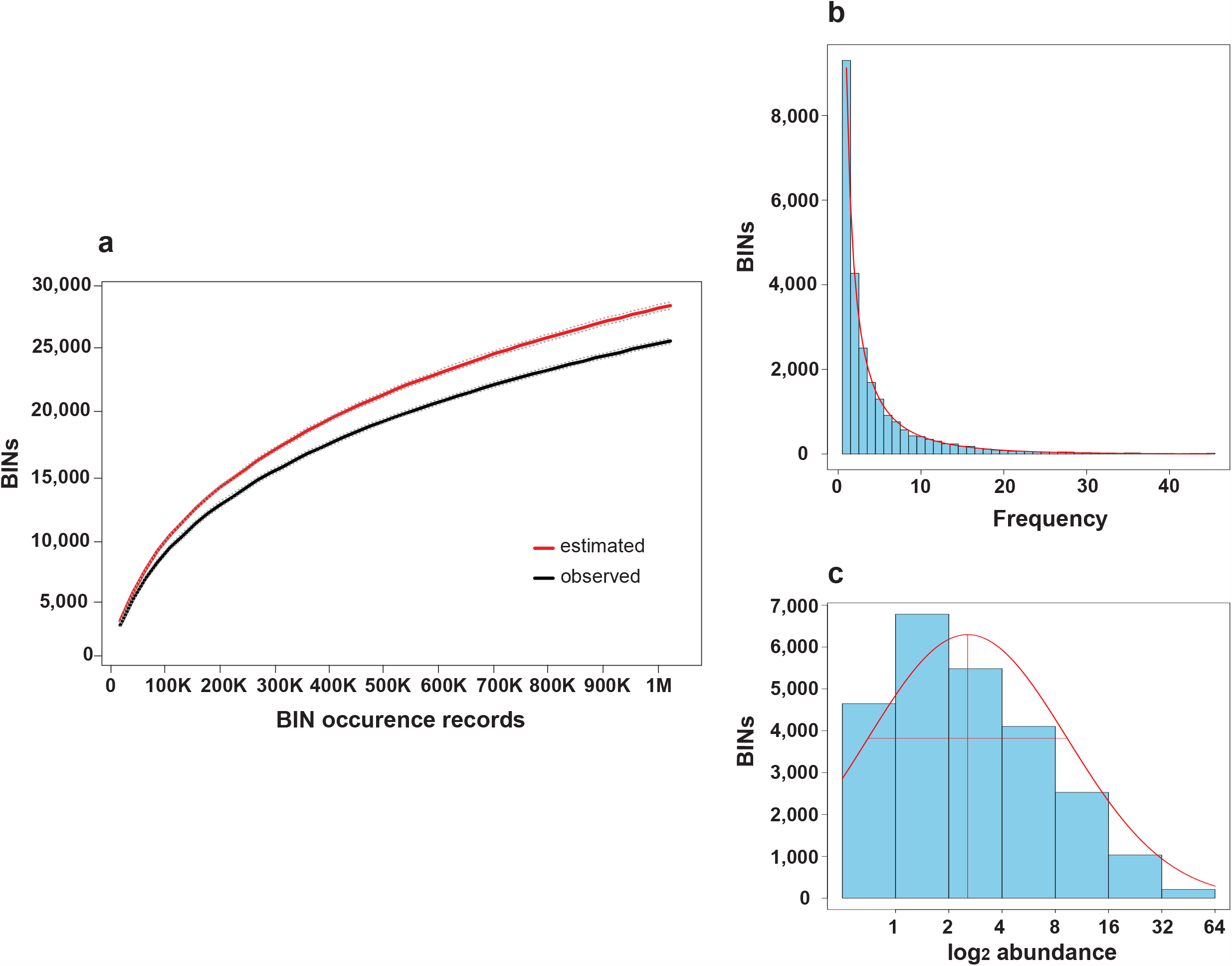

The Chao 1 [35] estimate for the total number of BINs present at the 52 sites was 29,640 (**Figure 2a**) while species richness extrapolation based on the lognormal distribution (**Figure 2c**, [36]) suggested the presence of 31,516 BINs. On average, 0.3 million sequences were recovered per sample, and they revealed the presence of an average of 2,352 BINs per site (range 996–4,581 BINs, **Table S2**) with bi-weekly samples containing an average of 619±14.3 S.E. BINs (range 60–1666, **Table S3**). Most low BIN counts occurred in spring (May) or fall (September) with diversity peaking in mid-summer (June/July). Taxonomic composition at an ordinal level was similar among samples with over half of the BINs being flies (Diptera), followed by Hymenoptera, Lepidoptera, Hemiptera, and Coleoptera.

Overlap in BIN composition was higher among parks in an ecoregion than among those in different ecoregions, even after geographical distance was considered (**Figure 3a**). Sites in the ECF had the highest mean phylogenetic diversity followed by EGL and finally SGL (**Figure 3b**), differences that were significant (KW and Dunn’s posthoc p < 0.003). More BINs were collected in the ECF (14,001) than in the EGL (12,787) or SGL (10,958) (Figure 3c). The Chao 1 estimates for the number of BINs present in each ecoregion were 15,401 for ECF, 14,577 for EGL, and 12,602 for SGL. The three ecoregions shared 4,133 BINs while about a third of those in each region were not collected elsewhere. A two-dimensional NMDS Ordination plot revealed that BIN assemblages for sites in each ecoregion formed cohesive groupings (Figure 3d). PERMANOVA analysis also suggested that community structure varied between ecoregions (*R*^2^ □=□ 0.141, *P*□ = □ 0.0001) and decreased site elevation (*R*^2^ □= □0.035, *P*□ = □0.03).

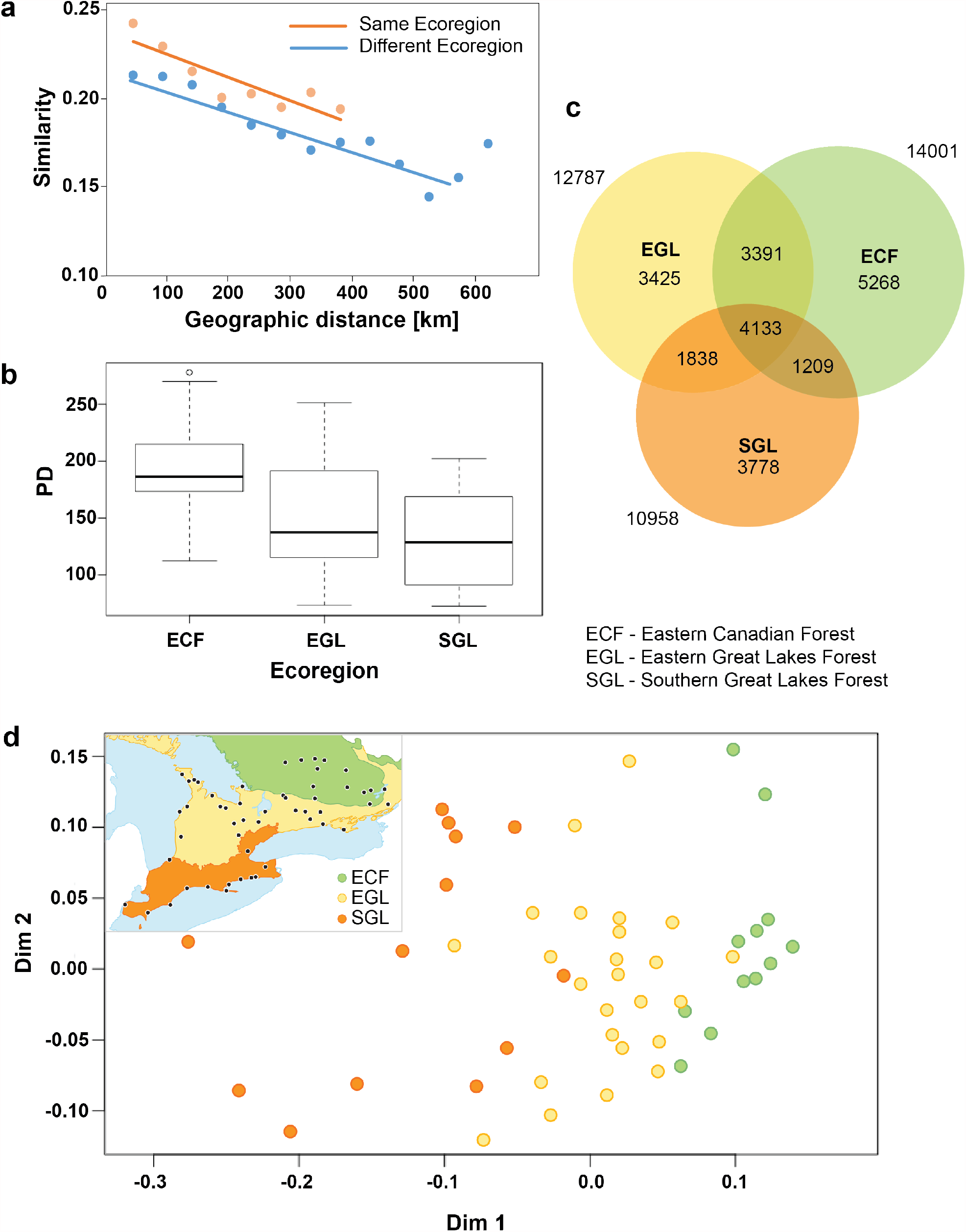

Overall, α-diversity was highest in the ECF, intermediate in the EGL, and lowest in SGL (**Figure 4**). The α-diversity patterns for the varied insect orders followed the overall trend, but BIN richness for Collembola showed the opposite trend as it peaked in the SGL, while spider α-diversity was highest in the EGL.

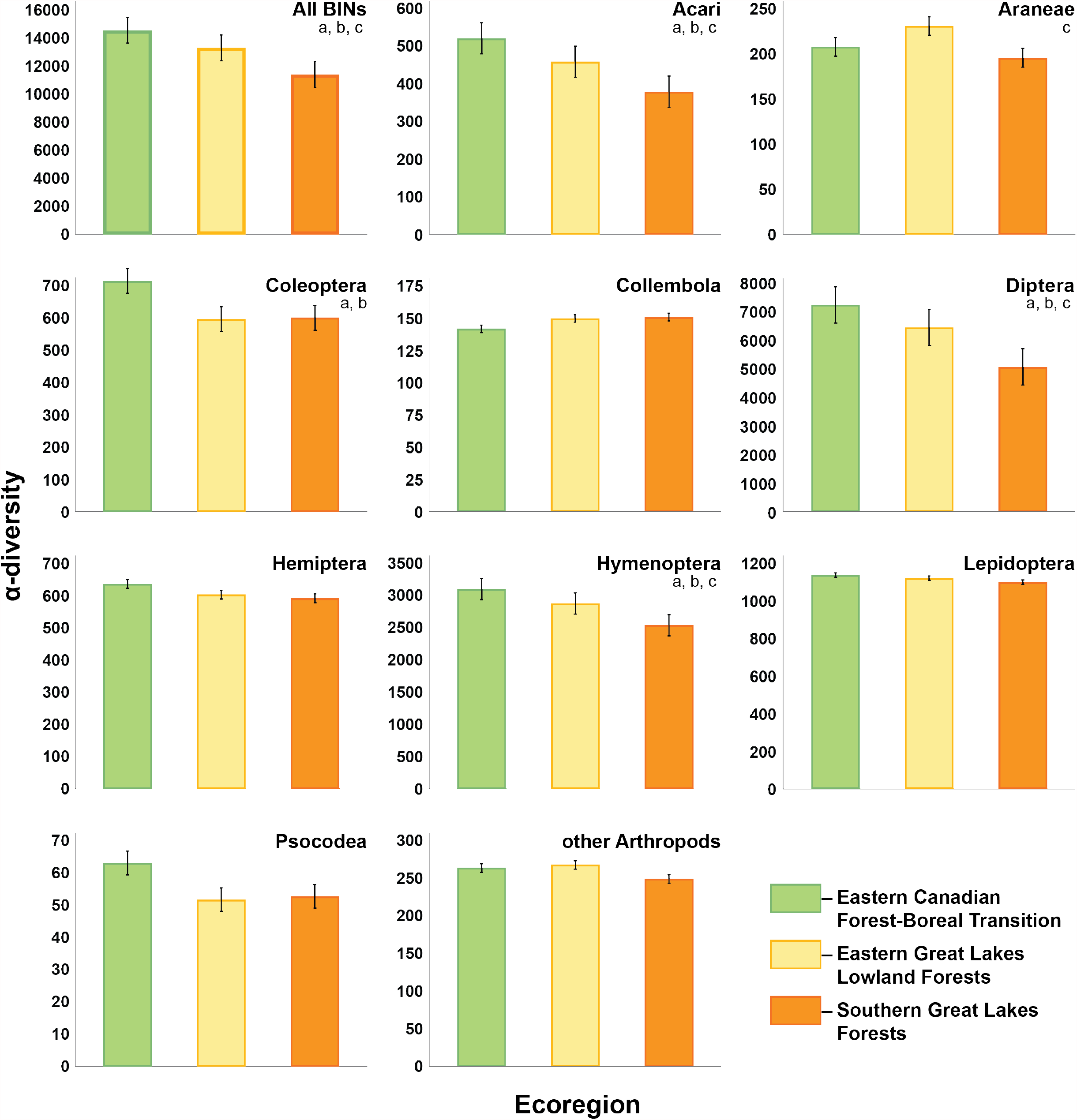

Levels of turnover (**Figure 5**) were generally high among sites (species replacement by new species not found elsewhere) as well as high nestedness levels (gain and loss of species also found elsewhere). Lower levels of both turnover and nestedness were observed for most taxa at sites in the ECF while the highest values were found in the SGL.

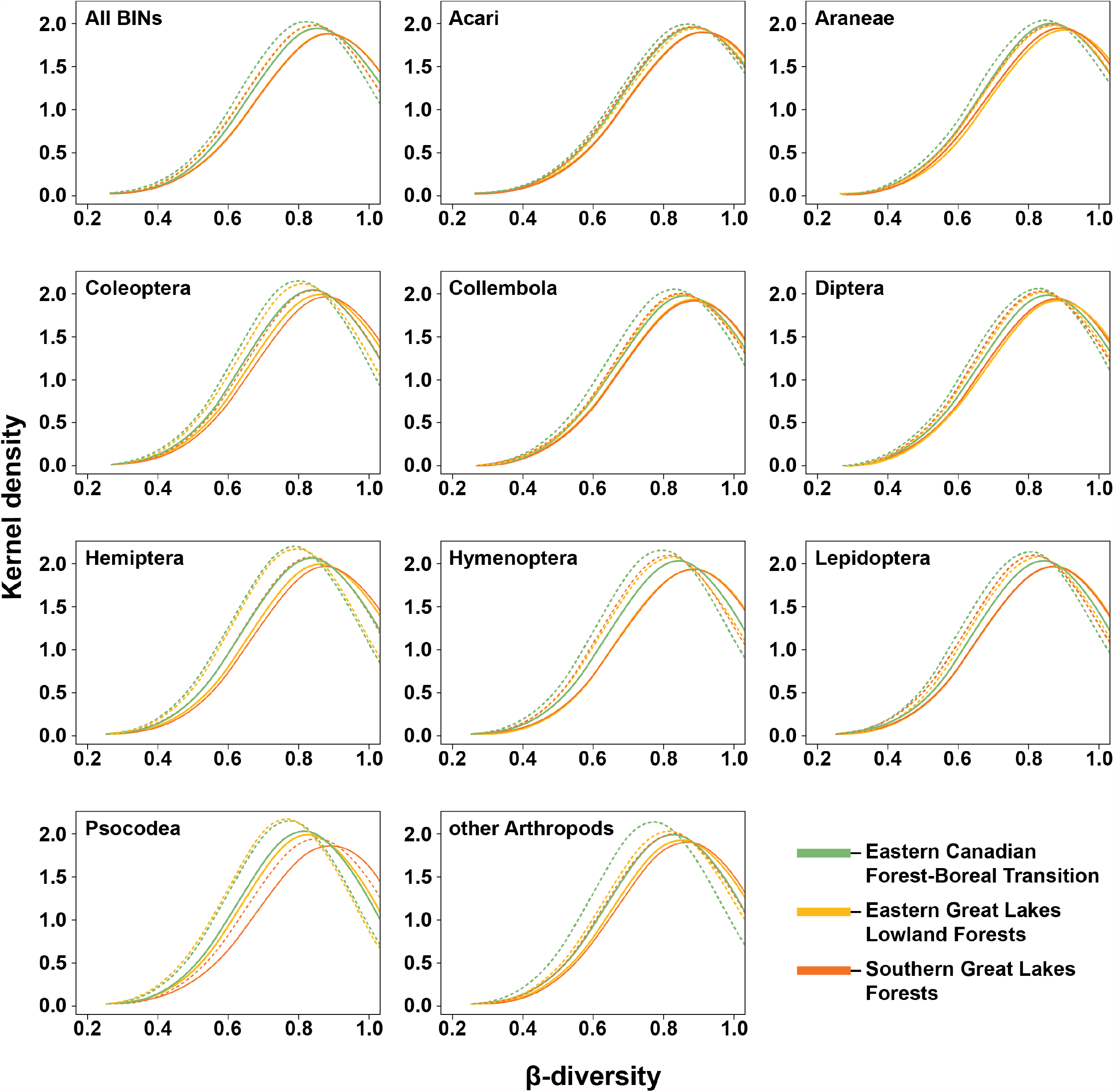

## Discussion

This study used metabarcoding to examine the species represented in 410 Malaise trap samples derived from 52 protected sites in three juxtaposed Canadian ecoregions. Metabarcoding revealed 26,263 species of arthropods while Chao 1 and Preston lognormal extrapolations indicated that another 3,000–5,000 species await detection. As just 52 sites were surveyed, a more comprehensive sampling program in these ecoregions might reveal as many as 50,000 species of arthropods. Nearly 5-fold variation (996– 4,581) in BIN counts were detected among sites; counts showed a similar range for the 30 sites where all samples were analyzed (996–4,508) and the 22 where just half were metabarcoded (1,312–4,581). On average, 619 BINs were recovered from each metabarcoded sample, a count that was 52.5% higher than the mean BIN count (406) for samples that were barcoded (Steinke et al. in prep). This difference suggests that more than half the BINs recovered from metabarcoded samples derive from environmental DNA attached to specimens in the sample or from their gut contents.

The three ecoregions examined in this study collectively span 160,000 km^2^, just 1.6% of Canada’s land surface, but two (SGL, EGL) are among the most heavily populated areas in the country [31]. The ecoregions showed considerable overlap in species composition; 33.1% of the BINs recorded from three or more sites were shared by the three ecoregions. BIN richness was lowest in the southernmost ecoregion (SGL) and highest in the most northerly (ECF). This difference coincided with a disturbance gradient -- from forested regions with low human density in the ECF (78% forest cover) to disturbed landscapes dominated by farmland/cities in the SGL (12% forest cover). The decline in species richness in response to disturbance is consistent with earlier studies [37-39], even though our collections all derived from protected areas. [40] reported that protected sites contain significantly higher species counts than adjacent disturbed areas, perhaps because communities in protected areas include representatives of original habitats and generalists from adjacent disturbed landscapes [41]. However, protected areas in the SGL were small islands of remnant forest in a landscape dominated by agricultural activity so they were undoubtedly heavily exposed to pesticides with agricultural fields creating dispersal barriers which further reduced diversity.

Our results indicate that α-diversity for major insect orders of flying insects (Diptera, Hymenoptera, Hemiptera, Lepidoptera) peaked in the least disturbed ecoregion (ECF). By contrast, two groups of arthropods (Araneae, Collembola) lacking flight showed a different trend with their diversity peaking in other ecoregions. This difference might reflect the fact that Malaise traps only sample flightless taxa with resident populations near the trap but capture flying insects from distant habitats. As such, biodiversity patterns for flying insects provide a regional perspective while those for taxa without flight provide a local perspective. If so, the reduction in diversity of Collembola from the most southerly (SGL) to northerly (ECF) ecoregion might reflect the expected latitudinal gradient in biodiversity, undisrupted by disturbance because of the local source of specimens in each sample.

The present study establishes the feasibility of monitoring temporal changes in species composition of arthropod communities [42,43]. For all three ecoregions, temporal turnover was high, reflecting the seasonal succession of species. β-diversity was lowest for most taxonomic groups at sites in the ECF and highest in the SGL. Species turnover was generally higher than nestedness, suggesting the presence of many transient species [44]. As many species were only collected at one or two sites, many samples likely included transients passively transported by the wind [45].

Metabarcoding can already provide cost-effective biosurveillance as the present study analyzed about 856,000 specimens and generated 223,860 species occurrence records for $82,000, an analytical cost of less than $0.50 per record. By adopting simpler analytical protocols (e.g., destructive processing of samples) with ongoing reductions in sequencing costs [10], costs can be reduced by an order of magnitude, delivering species occurrence records for $0.04 apiece in the ecoregions targeted in this study. In settings with higher α-diversity, the cost could be halved. Aside from its cost-effectiveness for data acquisition, the digital format of metabarcoding results aids their curation, validation, and preservation. Although current metabarcoding protocols cannot estimate the abundance of each species in a sample, the situation shifts when multiple samples are analyzed as the abundance of a species can then be estimated from its frequency of occurrence in these samples (rare species will be recovered less frequently than abundant taxa).

As the 846 currently recognized ecoregions [30] were largely delineated based on distributional data for vascular plants and vertebrates, there remains a need to ascertain how well they represent diversity patterns in other taxa. [46] found that arthropods showed weak adherence to ecoregion boundaries and proposed this might reflect dispersal limitations linked to their small body size or to the biased assemblage of arthropod species with data. Our much larger dataset shows evidence of structuring by ecoregion as both phylogenetic diversity and BIN composition were significantly different among ecoregions, even when comparisons extended to widely separated sites. This result suggests that ecoregions do provide a useful structural framework, reinforcing results from earlier studies [47,48]. However, a third of species in this study crossed ecoregion boundaries and more extensive sampling would raise the incidence of shared species. The latter results make it clear that high sampling effort is required to better understand species distributions. In looking to the future, it is apparent that there is an immediate need for a more detailed understanding of the levels of species overlap between adjacent ecoregions. Is, for example, the pattern of high overlap in species composition among neighbouring ecoregions detected in this study a general pattern or are some ecoregion boundaries sharply delineated? Such information is critical in designing an effective global biomonitoring network to inform conservation efforts [49,50].

## Potential Implications

Past monitoring programs have provided limited insights into the shifting distributions and abundances of arthropod species [51]. By coupling the use of an efficient collection method with the capacity of DNA metabarcoding to determine the species composition of bulk samples, this study has shown that compositional shifts in arthropod communities can be tracked [52]. The present results also indicate that the ecoregion concept not only furthers understanding of foundational biogeographic principles and improves their potential application to conservation efforts, but also provides a logical scaffold for large-scale monitoring networks.

## Methods

### Sample collection

An ez-Malaise trap (BioQuip Products) was deployed to collect arthropods at one site in each of 50 provincial parks while two sites were sampled in the final park (Algonquin) because of its large size. Trap catches were harvested every second week from early May through September, producing 10 samples per site for a total of 520 samples. These samples were preserved in 95% ethanol and held at -20° C until DNA extraction. Five samples (weeks 1+2, 5+6, 9+10, 13+14, 17+18) from each of 22 sites were employed for single specimen barcoding (Steinke et al., in prep) while the other 410 samples were analyzed in this study. A direct count indicated that 230,000 specimens were present in the 21.2% of the samples that were barcoded. On this basis, the remaining samples (78.8%), those examined in this study, included about 856,000 specimens.

### DNA extraction and PCR

DNA extraction employed a membrane-based protocol [32] modified for bulk samples. Specimens were removed from ethanol by filtration through a sterile Microfunnel 0.45 µM Supor Membrane Filter (Pall Laboratory) using a 6-Funnel Manifold (Pall Laboratory). The wet weight of each sample was then ascertained to allow volume adjustment (**Table S4**) of the lysis buffer [32]. Each sample was then incubated overnight at 56°C while gently mixed on a shaker. Eight 50 μl aliquots (technical replicates) from each of the 410 lysates were then transferred into 3,280 separate wells in 96-well microplates and DNA extracts were generated using Acroprep 3.0 µm glass fiber/0.2 µm Bio-Inert membrane plates (Pall Laboratory). Each plate contained 80 lysate samples, 8 technical replicates of a positive control (lysate from a bulk sample whose component specimens were individually Sanger sequenced – public BOLD dataset - dx.doi.org/10.5883/DS-AGAKS) and 8 negative controls. Each lysate was mixed with 100 μl of binding mix, transferred to a column plate, and centrifuged at 5000 g for 5 min. DNA was then purified with three washes; the first employed 180 μl of protein wash buffer centrifuged at 5000 g for 5 min. Each column was then washed twice with 600 μl of wash buffer centrifuged at 5000 g for 5 min. Columns were transferred to clean tubes and spun dry at 5000 g for 5 min to remove residual buffer before their transfer to clean collection tubes followed by incubation for 30 min at 56°C to dry the membrane. DNA was subsequently eluted by adding 60 μl of 10 mM Tris-HCl pH 8.0 followed by centrifugation at 5000 g for 5 min.

PCR reactions employed a standard protocol [53]. Briefly, each reaction included 5% trehalose (Fluka Analytical), 1× Platinum Taq reaction buffer (Invitrogen), 2.5 mM MgCl_2_ (Invitrogen), 0.1 μM of each primer (Integrated DNA Technologies), 50 μM of each dNTP (KAPA Biosystems), 0.3 units of Platinum Taq (Invitrogen), 2 μl of DNA extract, and Hyclone ultra-pure water (Thermo Scientific) for a final volume of 12.5 μl. Two-stage PCR was used to generate amplicon libraries for sequencing on an Ion Torrent S5 platform. The first round of PCR used the primer combination AncientLepF3 [54] and LepR1 [55] to amplify a 463 bp fragment of COI. Prior to the second PCR, first round products were diluted 2x with ddH_2_O. Fusion primers were then used to attach platform-specific unique molecular identifiers (UMIs) along with the sequencing adaptors required for Ion Torrent S5 libraries. Both rounds of PCR employed the same thermocycling conditions: initial denaturation at 94 °C for 2 min, followed by 20 cycles of denaturation at 94°C for 40 sec, annealing at 51°C for 1 min, and extension at 72 °C for 1 min, with a final extension at 72°C of 5 min.

### HTS library construction

For each plate, labelled products were pooled prior to sequencing. In total, 41 libraries were assembled. Each included eight technical replicates of 10 samples plus eight technical replicates of a negative and a positive control respectively (i.e., 96 samples). The ten samples from each of the 30 sites that were only metabarcoded, together with positive and negative controls, were pooled after UMI tagging to create a library that was analyzed on a 530 chip (30 chips in total). Five samples were available from each the other 22 sites (where half the samples were retained for barcoding). The UMI-tagged amplicons from five samples from each of two sites were pooled with positive and negative controls to produce a single library. Amplicon libraries were prepared on an Ion Chef (Thermo Fisher Scientific) following and sequenced on an Ion Torrent S5 platform at the Centre for Biodiversity Genomics following manufacturer’s instructions (Thermo Fisher Scientific).

### Sequence analysis

Reads from the eight replicates for each sample were concatenated using a bash script and uploaded to mBRAVE (http://mbrave.net/) for quality filtering and subsequent queries using several reference libraries in an open reference approach. All reads were queried against five system libraries on mBRAVE: bacteria (SYS-CRLBACTERIA), chordates (SYS-CRLCHORDATA), insects (SYS-CRLINSECTA), non-insect arthropods (SYS-CRLNONINSECTARTH), and non-arthropod invertebrates (SYS-CRLNONARTHINVERT). Sequences were only included in this analysis if they possessed a minimum length >350 bp and met the following three quality criteria (Mean QV >20; <25% positions with a QV<20; <5% positions with QV<10). Reads were trimmed 30 bp from their 5’ terminus with a set trim length of 450 bp. Reads were matched to the sequences in each reference library with an ID distance threshold of 3%, but were only retained for further analysis when at least three reads matched an OTU in the reference database. All reads failing to match any sequence in the five reference libraries were clustered at an OTU threshold of 1% with a minimum of five reads per cluster. All raw data are available in the NCBI Short Read Archive (PRJNA629553).

Using mBRAVE, we generated BIN (and OTU) tables including all library queries for each individual plate/run (10 samples, plus a negative and positive control - dx.doi.org/10.5883/DS-AGAKS - for each run). Read counts for any BINs recovered from the negative control on a plate were subtracted from the counts for the same BIN in the 80 non-control wells in the run. When this subtraction reduced the read count for a BIN to zero, its occurrence was removed. This step reduced the effects of rare tag switching on data integrity [56] and removed any background contamination.

### Ecoregion analysis

To determine the completeness of sampling, we calculated accumulation curves and the Chao-1 estimator for total diversity [35] using the vegan package [57]. For further extrapolation of species richness, we used the lognormal species abundance distribution [36]. The fit of Fisher’s Logseries [58] was used to determine relative BIN abundance. Both methods are implemented in vegan (fisherfit, prestonfit) [57]. We calculated Sørensen’s similarity coefficient to ascertain if differences in species assemblages were greater between or across ecoregion borders after controlling for distance. Differences in BIN composition among the three ecoregions were examined using non-metric multidimensional scaling (NMDS) with the Bray-Curtis index coefficient as implemented in vegan [57]. The adonis function of the vegan package was used to conduct a Permutational Multivariate Analysis of Variance (PERMANOVA) to partition distance matrices among sources of variation (factors such as latitude, longitude, elevation, and ecoregion).

A Maximum likelihood phylogeny was inferred for a BIN sequence alignment using RAxML Black box [59] on XCEDE via the CIPRES portal [60]. The resulting phylogeny comprising 26,263 BIN sequences was used to calculate Faith’s phylogenetic distance (PD) [61] using the picante package [62]. Because this measure is influenced by polytomies in a phylogeny [63], only one representative was included per BIN to avoid bias introduced by variation in the number of records for each BIN. A Kruskal-Wallis test followed by a Dunn’s posthoc analysis was used to determine if significant PD differences existed between ecoregions.

Alpha (α)-diversity was quantified as the number of BINs observed at a site. Beta (β)-diversity was computed as multi-site Sorensen and Simpson indices using the betapart 1.3. package [64]. β-diversity calculations between pairs of ecoregions were computed using 12 random sites from the total sites for each ecoregion, and resampled 1000 times. We then decomposed the among-site β-diversity into its turnover (species replacement from site to site) and nestedness (species gain/loss from sites) components. Pairwise BIN diversity among ecoregions was evaluated using the nonparametric multiple comparison function implemented in the R package dunn.test 1.2.4 [65]. dunn.test is equivalent to the Kruskall–Wallis and pair-wise Mann–Whitney post hoc tests with Bonferroni correction.

All analyses were performed in R v.3.4.4 [66].

## Supporting information

Suppmental Table 3

## Author contributions

DS, EVZ, JRDW, PDNH designed the study. DS, JRDW, JES, KP coordinated the study. SLDW, NVI, SWJP, TWAB did the bench work and contributed to analyses. SR and MM oversaw database organisation. DS did the analyses and wrote the manuscript. PDNH, JRDW, EVZ, TWAB revised the manuscript.

## Acknowledgements

We thank the collections and sequencing staff at the Centre for Biodiversity Genomics for acquiring and processing the specimens analyzed in this study. We are very grateful to Suz Bateson for improving the figures and to staff at the participating Ontario Provincial Parks for facilitating collections.

