## Supplementary figures and images for "Message in a Bottle – Metabarcoding Enables Biodiversity Comparisons Across Ecoregions"

### Supplemental Figure 1

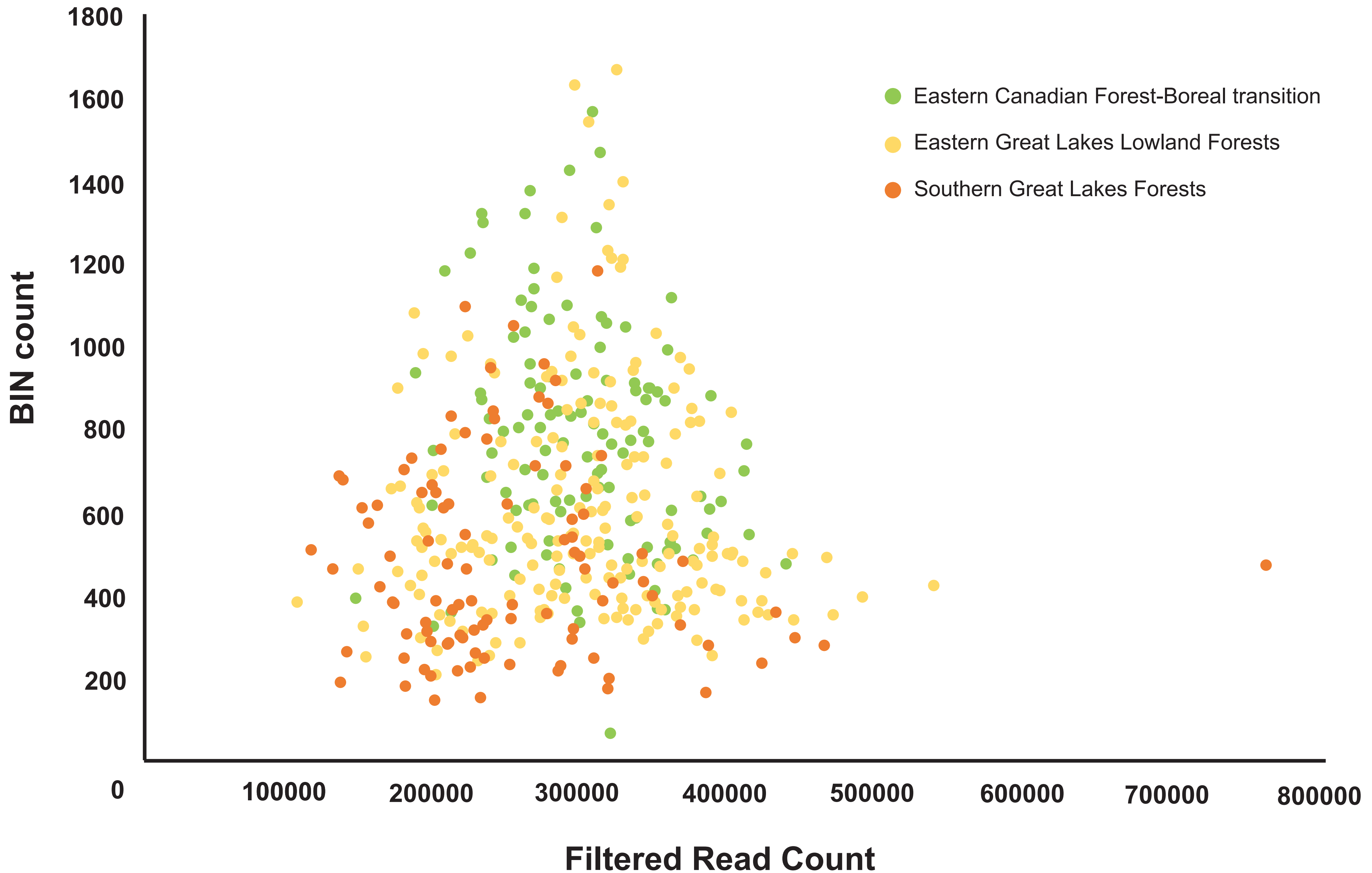
