## Supplemental Table 1 for "Message in a Bottle – Metabarcoding Enables Biodiversity Comparisons Across Ecoregions"

**Table S1:** mBRAVE project codes as well as samples analyzed and read coverage for each 530 chip analyzed on the Ion Torrent S5. The 11 chips that included samples from two sites are highlighted in yellow.

| **Project Code** | **Project Title** | **Reads** | **Filtered Reads** |
| --- | --- | --- | --- |
| MBR-OPPMAA | Ontario Provincial Parks 2014:CCDB-S5-0053_CBGMB-00003_Algonquin PP - Rock Lake Site 1 | 9376304 | 2592428 |
| MBR-OPPMABMAC | Ontario Provincial Parks 2014:CCDB-S5-0066_CBGMB-00016_Algonquin PP - Oxtongue Site 3 + Awenda PP | 8544988 | 2674401 |
| MBR-OPPMADMAE | Ontario Provincial Parks 2014:CCDB-S5-0067_CBGMB-00017_Balsam Lake PP + Bayview Escarpment PP | 8535733 | 3294349 |
| MBR-OPPMAFMAI | Ontario Provincial Parks 2014:CCDB-S5-0068_CBGMB-00018_Bell Bay PP + Boyne Valley PP | 6430734 | 1847104 |
| MBR-OPPMAHMBD | Ontario Provincial Parks 2014:CCDB-S5-0086_CBGMB-00035_Bon Echo PP - Site 1 + Morris Tract PP | 7271497 | 2424720 |
| MBR-OPPMAJ | Ontario Provincial Parks 2014:CCDB-S5-0057_CBGMB-00007_Bronte Creek PP | 6892296 | 2517868 |
| MBR-OPPMAKMAO | Ontario Provincial Parks 2014:CCDB-S5-0073_CBGMB-00019_Charleston Lake PP + Ferris PP | 10020289 | 3480857 |
| MBR-OPPMAL | Ontario Provincial Parks 2014:CCDB-S5-0058_CBGMB-00008_Duncan Escarpment PP | 9293842 | 3435885 |
| MBR-OPPMAN | Ontario Provincial Parks 2014:CCDB-S5-0061_CBGMB-00009_Emily PP | 8315694 | 3169017 |
| MBR-OPPMAP | Ontario Provincial Parks 2014:CCDB-S5-0063_CBGMB-00010_Forks of the Credit PP | 10348276 | 4090743 |
| MBR-OPPMAQMAU | Ontario Provincial Parks 2014:CCDB-S5-0069_CBGMB-00020_Frontenac PP + Inverhuron PP | 7931246 | 2871585 |
| MBR-OPPMAR | Ontario Provincial Parks 2014:CCDB-S5-0064_CBGMB-00012_Holland Landing Prairie PP | 8124964 | 2701838 |
| MBR-OPPMAS | Ontario Provincial Parks 2014:CCDB-S5-0054_CBGMB-00004_Hope Bay Forest PP | 9227593 | 3463105 |
| MBR-OPPMAT | Ontario Provincial Parks 2014:CCDB-S5-0065_CBGMB-00013_Indian Point PP - Site 1 | 9151934 | 3599629 |
| MBR-OPPMAWMAZ | Ontario Provincial Parks 2014:CCDB-S5-0070_CBGMB-00021_John E Pearce PP + Lions Head PP | 10693129 | 3641703 |
| MBR-OPPMAX | Ontario Provincial Parks 2014:CCDB-S5-0080_CBGMB-00026_Johnston Harbour PP | 7927583 | 2339035 |
| MBR-OPPMAY | Ontario Provincial Parks 2014:CCDB-S5-0081_CBGMB-00027_Lake St Peter PP | 9380178 | 3266285 |
| MBR-OPPMBAMBE | Ontario Provincial Parks 2014:CCDB-S5-0072_CBGMB-00022_Lower Madawaska River PP + Murphys Point PP | 10315593 | 3815438 |
| MBR-OPPMBB | Ontario Provincial Parks 2014:CCDB-S5-0082_CBGMB-00028_MacGregor Point PP | 9944565 | 3854418 |
| MBR-OPPMBC | Ontario Provincial Parks 2014:CCDB-S5-0056_CBGMB-00006_Mark S Burnham PP | 10796469 | 4536699 |
| MBR-OPPMBFMBH | Ontario Provincial Parks 2014:CCDB-S5-0074_CBGMB-00023_Ojibway Prairie PP + Pinery PP - Site 2 | 9804179 | 3243861 |
| MBR-OPPMBG | Ontario Provincial Parks 2014:CCDB-S5-0083_CBGMB-00029_Petroglyphs PP | 10645785 | 3989178 |
| MBR-OPPMBI | Ontario Provincial Parks 2014:CCDB-S5-0084_CBGMB-00030_Port Burwell PP | 10140496 | 3058298 |
| MBR-OPPMBJ | Ontario Provincial Parks 2014:CCDB-S5-0085_CBGMB-00031_Presqu'ile PP | 10651981 | 4259741 |
| MBR-OPPMBMMBN | Ontario Provincial Parks 2014:CCDB-S5-0075_CBGMB-00024_Rondeau PP - Site 1 + Sandbanks PP | 10725061 | 3868406 |
| MBR-OPPMBO | Ontario Provincial Parks 2014:CCDB-S5-0087_CBGMB-00036_Selkirk PP | 9153371 | 2649363 |
| MBR-OPPMBP | Ontario Provincial Parks 2014:CCDB-S5-0055_CBGMB-00005_Sharbot Lake PP | 8344274 | 3368635 |
| MBR-OPPMBQMBU | Ontario Provincial Parks 2014:CCDB-S5-0079_CBGMB-00025_Short Hills PP + Turkey Point PP | 7941400 | 2702280 |
| MBR-OPPMBR | Ontario Provincial Parks 2014:CCDB-S5-0088_CBGMB-00037_Sibbald Point PP | 8752087 | 3151909 |
| MBR-OPPMBS | Ontario Provincial Parks 2014:CCDB-S5-0089_CBGMB-00038_Silent Lake PP | 8381273 | 2870078 |
| MBR-OPPMBT | Ontario Provincial Parks 2014:CCDB-S5-0090_CBGMB-00039_Silver Lake PP | 8105235 | 2674401 |
| MBR-OPPMBX | Ontario Provincial Parks 2014:CCDB-S5-0097_CBGMB-00040_Wheatley PP | 9036296 | 2592428 |
| MBR-OPPMBZZ | Ontario Provincial Parks 2014:CCDB-S5-0052_CBGMB-00002_Peter's Woods PP Malaise | 8048432 | 3302341 |
| MBR-OPPMBY | Ontario Provincial Parks 2014:CCDB-S5-041_CBGMB-00001_Long Point PP Malaise | 11177387 | 4232828 |
| MBR-OPPMAG | Ontario Provincial Parks 2014:CCDB-S5-0127_CBGMB-00068_Black Creek PP | 7877916 | 2491752 |
| MBR-OPPMAM | Ontario Provincial Parks 2014:CCDB-S5-0128_CBGMB-00069_Earl Rowe PP | 8955778 | 3043981 |
| MBR-OPPMAV | Ontario Provincial Parks 2014:CCDB-S5-0129_CBGMB-00070_James N Allan PP | 6594231 | 1605985 |
| MBR-OPPMBK | Ontario Provincial Parks 2014:CCDB-S5-0130_CBGMB-00071_Pretty River Valley PP | 7497170 | 2158049 |
| MBR-OPPMBL | Ontario Provincial Parks 2014:CCDB-S5-0131_CBGMB-00072_Rock Point PP | 8479191 | 1624386 |
| MBR-OPPMBV | Ontario Provincial Parks 2014:CCDB-S5-0132_CBGMB-00073_Upper Madawaska River PP | 9938090 | 3074949 |
| MBR-OPPMBW | Ontario Provincial Parks 2014:CCDB-S5-0133_CBGMB-00074_Wasaga Beach PP | 9050667 | 2673304 |
