## Supplemental Table 2 for "Message in a Bottle – Metabarcoding Enables Biodiversity Comparisons Across Ecoregions"

**Table S2:** GPS coordinates, elevation (m), and ecoregion assignment for the 52 sampling sites and the number of BINs recovered from each site. ECF = Eastern Canadian Forest (15 sites); EGL = Eastern Great Lakes Forests (24 sites); SGL = Southern Great Lakes Forests (13 sites).

| **Provincial Park** | **Latitude** | **Longitude** | **Elevation(m)** | **BINs** | **Ecoregion** |
| --- | --- | --- | --- | --- | --- |
| Algonquin - Oxtongue River | 45.4621 | -78.796 | 427 | 1642 | ECF |
| Algonquin - Rock Lake | 45.51962 | -78.39752 | 353 | 2030 | ECF |
| Bell Bay | 45.51506 | -77.81812 | 320 | 3186 | ECF |
| Bon Echo | 44.89405 | -77.19691 | 272 | 2638 | ECF |
| Charleston Lake | 44.49798 | -76.0414 | 88 | 3021 | ECF |
| Frontenac | 44.51783 | -76.53944 | 166 | 3498 | ECF |
| Indian Point | 44.60414 | -78.82895 | 258 | 2403 | ECF |
| Lake St Peter | 45.3202 | -78.02496 | 406 | 2664 | ECF |
| Lower Madawaska River | 45.25606 | -77.19221 | 262 | 3333 | ECF |
| Murphys Point | 44.78118 | -76.2336 | 132 | 4581 | ECF |
| Petroglyphs | 44.61605 | -78.04084 | 265 | 3194 | ECF |
| Sharbot Lake | 44.77952 | -76.72379 | 201 | 2878 | ECF |
| Silent Lake | 44.92144 | -78.06931 | 360 | 2895 | ECF |
| Silver Lake | 44.83129 | -76.57565 | 184 | 4508 | ECF |
| Upper Madawaska River | 45.53535 | -78.04699 | 318 | 3796 | ECF |
| Awenda | 44.82534 | -79.98458 | 224 | 3135 | EGL |
| Balsam Lake | 44.62857 | -78.8614 | 274 | 3227 | EGL |
| Bayview Escarpment | 44.63367 | -80.69829 | 329 | 3966 | EGL |
| Black Creek | 44.96797 | -81.36156 | 179 | 2369 | EGL |
| Boyne Valley | 44.11563 | -80.12777 | 460 | 1609 | EGL |
| Duncan Escarpment | 44.42305 | -80.46923 | 395 | 1957 | EGL |
| Earl Rowe | 44.15176 | -79.903 | 216 | 1479 | EGL |
| Emily | 44.34143 | -78.53746 | 251 | 1916 | EGL |
| Ferris | 44.28286 | -77.79627 | 132 | 4005 | EGL |
| Forks of the Credit | 43.82415 | -80.00309 | 403 | 2753 | EGL |
| Holland Landing Prairie | 44.11894 | -79.48795 | 225 | 2545 | EGL |
| Hope Bay Forest | 44.92509 | -81.15563 | 253 | 1071 | EGL |
| Inverhuron | 44.29838 | -81.59065 | 182 | 1785 | EGL |
| Johnston Harbour-Pine Tree Point | 45.1171 | -81.53679 | 177 | 1939 | EGL |
| Lions Head | 44.99539 | -81.2334 | 219 | 2074 | EGL |
| MacGregor Point | 44.41072 | -81.44641 | 191 | 2607 | EGL |
| Mark S Burnham | 44.29882 | -78.26779 | 209 | 1633 | EGL |
| Morris Tract | 43.72995 | -81.64166 | 257 | 1673 | EGL |
| Peters Woods | 44.12845 | -78.04057 | 236 | 4197 | EGL |
| Presqu'ile | 44.00914 | -77.7424 | 77 | 3197 | EGL |
| Pretty River Valley | 44.41232 | -80.30035 | 337 | 3550 | EGL |
| Sandbanks | 43.90287 | -77.26929 | 85 | 1312 | EGL |
| Sibbald Point | 44.32982 | -79.32737 | 221 | 1872 | EGL |
| Wasaga Beach | 44.51258 | -80.01165 | 188 | 1314 | EGL |
| Bronte Creek | 43.4023 | -79.7617 | 125 | 2800 | SGL |
| James N Allan | 42.84962 | -79.66397 | 175 | 2029 | SGL |
| John E Pearce | 42.60595 | -81.44243 | 195 | 1339 | SGL |
| Long Point | 42.58006 | -80.38538 | 175 | 1363 | SCF |
| Ojibway Prairie | 42.26278 | -83.07246 | 180 | 3311 | SGL |
| Pinery | 43.26987 | -81.82706 | 182 | 1978 | SGL |
| Port Burwell | 42.6544 | -80.81493 | 196 | 996 | SGL |
| Rock Point | 42.85405 | -79.55536 | 176 | 2246 | SGL |
| Rondeau | 42.30206 | -81.85306 | 175 | 1356 | SGL |
| Selkirk | 42.81676 | -79.95736 | 180 | 1146 | SGL |
| Short Hills | 43.11288 | -79.27376 | 95 | 2541 | SGL |
| Turkey Point | 42.70515 | -80.32849 | 222 | 2513 | SGL |
| Wheatley | 42.09199 | -82.44227 | 181 | 1210 | SGL |
