## Supplemental Table 4 for "Message in a Bottle – Metabarcoding Enables Biodiversity Comparisons Across Ecoregions"

**Table S4:** Wet weight (g) to insect lysis buffer volume (mL) ratios for Malaise trap bulk samples.

| **Wet Weight of Bulk Sample (g)** | **Insect Lysis Buffer Volume (mL)** |
| --- | --- |
| <1.5 | 15 |
| 1.5-4.9 | 20 |
| 5.0-9.9 | 50 |
| 10.0-19.9 | 100 |
| 20.0-29.9 | 200 |
| >30.0 | 250 |
